# Possible biased virulence attenuation in the Senegal strain of *Ehrlichia ruminantium* by *ntrX* gene conversion from an inverted segmental duplication

**DOI:** 10.1101/2020.11.26.400648

**Authors:** Jonathan L. Gordon, Adela S. Oliva Chavez, Dominique Martinez, Nathalie Vachiery, Damien F. Meyer

## Abstract

*Ehrlichia ruminantium* is a tick-borne intracellular pathogen of ruminants that causes heartwater, a disease present in Sub-saharan Africa, islands in the Indian Ocean and the Caribbean, inducing significant economic losses. At present, three avirulent strains of *E. ruminantium* (Gardel, Welgevonden and Senegal isolates) have been produced by a process of serial passaging in mammalian cells *in vitro*, but unfortunately their use as vaccines do not offer a large range of protection against other strains, possibly due to the genetic diversity present within the species (Cangi et al. 2016). So far no genetic basis for virulence attenuation has been identified in any *E. ruminantium* strain that could offer targets to facilitate vaccine production. Virulence attenuated Senegal strains have been produced twice independently, and require many fewer passages to attenuate than the other strains. We compared the genomes of a virulent and attenuated Senegal strain and identified a likely attenuator gene, *ntrX*, a global transcription regulator and member of a two-component system that is linked to environmental sensing. This gene has an inverted partial duplicate close to the parental gene that shows evidence of gene conversion in different *E. ruminantium* strains. The pseudogenisation of the gene in the avirulent Senegal strain occurred by gene conversion from the duplicate to the parent, transferring a 4bp deletion which is unique to the Senegal strain partial duplicate amongst the wild isolates. We confirmed that the *ntrX* gene is not expressed in the avirulent Senegal strain by RT-PCR. The inverted duplicate structure combined with the 4bp deletion in the Senegal strain can explain both the attenuation and the faster speed of attenuation in the Senegal strain relative to other strains of *E. ruminantium*. Our results identify *nrtX* as a promising target for the generation of attenuated strains of *E. ruminantium* by random or directed mutagenesis that could be used for vaccine production.

## Introduction

*Ehrlichia ruminantium*, the causative agent of heartwater, is an obligate intracellular Gram-negative α-proteobacterium of the order *Rickettsiales*. It is a pathogen of domestic and wild ruminants transmitted by ticks from the *Amblyomma* genus. Heartwater is present in sub-Saharan Africa, the Indian Ocean and the Caribbean, and is an invasion threat for the Americas with the potential spread of *E. ruminantium-infected* ticks by migratory birds (Vachiéry et al. 2013). Furthermore, Heartwater is considered to be one of the 12 priority transboundary animal diseases by the US Homeland security department for the American mainland (Roth, Richt, and Morozov 2013). In the mammalian host, *E. ruminantium* principally infects endothelial cells, mainly from brain capillarties, but has been shown to also infect macrophages (Plessis 1975) and neutrophils (Camus and Barré 1988). *E. ruminantium* causes up to 82% mortality in susceptible ruminants (Allsopp 2010), and in endemic areas the economic importance of this pathogen is comparable to trypanosomiasis and East Coast Fever (Allsopp 2010). The economic importance of the disease in endemic areas and its potential risk of spreading into mainland North America have increased the interest in the development of a vaccine (Vachiéry et al. 2013), however, due to a large genetic variability between strains it is difficult to develop effective vaccines that offer cross-protection against multiple strains (Allsopp 2010; Zweygarth et al. 2005).

Like other members of the Anaplasmataceae family, *E. ruminantium* possesses a biphasic life cycle presenting two morphologically different forms that can be identified by electron microscopy (Jongejan et al. 1991; Bell-Sakyi et al. 2000). The dense-core or elementary body (EB) is the non-dividing infectious form, and the reticulate body (RB) is the replicative form. Both forms are present in the vertebrate host and the tick vector. Initially the EB attaches to the host endothelial cell and enters by endocytosis, developing a safe replicative niche where it differenciates into RB form that divides to form large colonies called morulae. After a few days, RB redifferenciate into infectious EB that are released from infected host cells by complete cell lysis and initiate a new infectious cycle (Moumène and Meyer 2016). Although not much is known about the differences in the gene or protein expression of these two forms in *E. ruminantium*, other members of the Anaplasmataceae family have been extensively studied, showing essentially different profiles of expression between the two forms (Mastronunzio, Kurscheid, and Fikrig 2012). Furthermore, form-specific markers have been identified in two members of the Anaplasmataceae families, *Anaplasma phagocytophilum* (Troese et al. 2011) and *Ehrlichia chaffeensis* (Popov, Yu, and Walker 2000).

Attenuation of pathogen virulence through *in vitro* passage is a common practice, however, in the case of *Rickettsiales*, only a few cases of attenuation have been reported (Jongejan 1991; Jongejan et al. 1993; Or et al. 2009), and the possible mechanism of attenuation has only been studied in limited cases (Bechah et al. 2010). Detailed studies of the mechanism of attenuation by genomic comparison of two *Rickettsia prowazekii* Madrid E vaccine strains (attenuated) versus Evir (reverted virulent strain) showed that the mutation of a single gene (*smt*) encoding a SAM-methyltransferase led to the attenuation of the bacteria (Y. Liu et al. 2014). The authors identified the only similarity between the two attenuated strains (DHW and DOW) was the frameshift mutation of *smt* in the attenuated strain (Y. Liu et al. 2014). Often mutation of global regulators can cause attenuation because a small perturbation at the top of a regulation network can lead to large and widespread pleiotropic changes in gene expression (Parkinson and Kofoid 1992; Cheng et al. 2006; Nene and Kole 2008; Jansen et al. 2015).

In an attempt to produce vaccines, virulence attenuated strains from Guadeloupe (Gardel), South Africa (Welgevonden) and Senegal have been generated *in vitro* by passaging the bacterium in bovine endothelial cells (and canine macrophage-monocyte cells for Welgevonden)(Jongejan 1991; Zweygarth et al. 2005; Pilet et al. 2012; Marcelino et al. 2015). The number of passages until attenuation differs largely between the strains, with Senegal having taken only 11 passages and less than 20 passages in two independent attempts to attenuate compared to 230 passages for Gardel in bovine endothelial cells (Jongejan 1991; Zweygarth et al. 2005; Pilet et al. 2012; Marcelino et al. 2015). Welgevonden strain on the other hand was attenuated by passage 56 in canine macrophage-monocytes and then readapted to bovine endothelial cells and failed to attenuate by passage 231 in bovine endothelial cells alone (Zweygarth et al. 2005). Usage of vaccines in the field has remained limited due to constraints associated with storage of live vaccines as they require liquid nitrogen. Moreover, attenuated vaccines, as for any other vaccine against heartwater, protect against homologous challenges but confer limited protection against heterologous strains, which is problematic as co-occurrence of different strains in infected animals is common (Zweygarth et al. 2005; Frutos et al. 2006). Understanding the mechanisms that lead to the attenuation of these particular strains may lead to the discovery of targets to generate more effective vaccines. So far however, there has been no genetic explanation for the attenuation of any of the *E. ruminantium* avirulent stocks.

In order to better understand the attenuation process in the Senegal strain of *Ehrlichia ruminantium*, we sequenced the genome of the virulent strain (ERSA) and an attenuated strain (ERSB) and compared their genomes to find molecular differences that differentiate them. Within the set of genomic differences there should be at least one that can explain the attenuation process in the Senegal strain and potentially why it has occured in far fewer *in vitro* passages in the Senegal strain than the other *Ehrlichia* strains. Here we present the results of this analysis, highlighting the *ntrX* gene, which is disrupted by a process of gene conversion from a nearby inverted partial duplication. The *ntrX* gene codes for the response regulator of the NtrY/NtrX two component system (2CS), one of only three such systems found in Ehrlichia species (Cheng et al. 2006; Cheng, Lin, and Rikihisa 2014; Rikihisa 2015), which are generally involved in the sensing of environmental or cellular signals and the coordination of the regulation of an array of genes in response to these signals. This study represents the first probable genetic explanation of virulence attenuation in *Ehrlichia ruminantium*.

## Methods

### DNA extraction and purification

The virulent and attenuated strains are those described in (Pilet et al. 2012). *Ehrlichia ruminantium* Senegal strain passage 7 (80% lysis at day 21 post infection (pi)) and 63 and 64 (80% lysis at day 7 pi) were cultured in Bovine Aortic endothelial cells as previously described for Gardel strain (Marcelino et al. 2005). When 80% lysis had occurred, supernatant and cellular debris were collected and centrifuged for 15 minutes at 4,000g at 4°C to remove cellular debris. The supernatant was then centrifuged for 30 minutes at 20,000g at 4°C in order to collect elementary bodies and remove the supernatant. The pellet was resuspended in 2ml of cold PBS and homogenized gently then 20 ml of PBS was added to wash it, followed by centrifugation for 30 minutes at 20,000g at 4°C. The pellet was resuspended in 350μl PBS and 10μl of RNase at 10mg/ml (SIGMA, Lyon, France) and 150μl of DNase I (Roche, Boulogne-Billancourt, France) were added. The pellet was incubated at 37°C for 90 minutes and the reaction was stopped by adding 25μl 0.5M EDTA, pH8. The DNase and Rnase were removed by centrifugation for 15 minutes at 20,000g at 12°C followed by a wash with 900μl of sterile DNase and RNase free water. Cells were centrifuged under the same conditions and the washing step was repeated twice. Lysis of elementary bodies was performed by adding 500μl of lysis solution (0.1M TRIS-HCl (pH8), 0.15M NaCl, 0.025M EDTA (pH8), 1.5% SDS, 0.3 mg/ml Proteinase K) followed by Incubation for 120 minutes at 55°C. DNA extraction was performed using phenol/chloroform (Perez et al. 1997) as follows: 500μl of phenol (Eurobio, Courtaboeuf, France) was added to 500μl of sample and mixed gently to homogenize followed by centrifugation for 5 minutes at 8000 g. The aqueous phase was collected and an equivalent volume of phenol was added before centrifugation for 5 minutes at 8000 g. The aqueous phase was collected and an equivalent volume of phenol (Eurobiotech, France), chloroform and isoamylalcool (Prolabo, Normapur, France) in 24:24:1 proportion) was added and mixed gently. The mixture was centrifuged for 5 minutes at 8000g and the aqueous phase collected and mixed gently with an equal volume of chroroform/isoamylalcool in 48:1 proportion. The mixture was centrifuged for 5 minutes at 8000g and the supernatant was collected and precipitated in ethanol (Prolabo, Normapur) by adding 2 volumes of absolute ethanol for 1 volume of the collected sample. The sample was stored at 4°C for one hour in order to obtain a white precipitate which was then centrifuged for 10 minutes at 10000g at 10°C. The supernatant was removed and the pellet resuspended in 1 ml of 75% ethanol followed by centrifugation for 5 minutes at 10000g. The supernatant was removed and the pellet air dried. The pellet was resuspended in 25μl of TE buffer (10mM TRISpH8, 1mM EDTA pH8). DNAs from Senegal passage 7, 63 and 64 in TE buffer were stored at −20°C before being used for further sequencing.

### Genome sequencing and assembly

The genome of the virulent Senegal strain (passage 7) was sequenced using 454 GS FLX technology. The attenuated strain (passage 63 & 64) was sequenced using 454 GS FLX technology with additional Sanger sequencing. Adapters for the 454 sequences were clipped with Cutadapt (Martin 2011), and the 454 reads were error corrected with Fiona (Schulz et al. 2014). The virulent strain was assembled using Mira 4.0.2 (Chevreux, Wetter, and Suhai 1999), resulting in 8 contigs larger than 1kb. The contigs were ordered according to the previously published Welgevonden strain genome (Collins et al. 2005) using Mauve (A. E. Darling, Mau, and Perna 2010). The attenuated strain reads were mapped onto the assembled virulent strain by using Bowtie 2 (Langmead and Salzberg 2012). Duplicate reads were removed using Picard (“Picard Tools” 2016), conversion between SAM and BAM formats, sorting and mpileup was done using Samtools (Li et al. 2009) and variants were called using BCFtools (Li et al. 2009). Indels and SNPs were identified with quality cutoffs of 50 and 20 respectively and were manually checked by viewing in Tablet (Milne et al. 2013). The virulent strain assembly was then edited to produce the attenuated genome. Both genome assemblies were submitted to Genbank (Accessions XX,YY).

### *E. ruminantium* RNA purification for RT-PCR

*E. ruminantium* Senegal attenuated (passage 63 and 66) and virulent (passage 7 and 11) were inoculated in 25 cm^2^ tissue culture flask containing bovine aortic endothelial cells as previously described in (Marcelino et al. 2005). The medium was changed at 24h and 72h for the attenuated strain and at 24h and every two days, thereafter, for the virulent strain. Cell layers were allowed to reach around 80-90% lysis and were mechanically harvested with a scraper. Re-suspended cells were centrifuged at 4,500 x g for 30 minutes at 4 °C. Supernatant was discarded and cells were re-suspended in 1 ml phosphate-buffered saline (PBS). Cells were then centrifuged at 10000 x g for 10 minutes and the PBS was removed. Pellets were stored at −70°C until RNA purification. Cell pellets were allowed to thaw for 5 minutes in ice and RNA was purified using the SV total RNA isolation system (Promega Corporation, Wisconsin, USA). An additional DNAse treatment was added by using the rigorous DNAse treatment with Turbo DNA-free (Ambion, Fisher Scientific, Illkirch, France), which consisted in adding 0.5 ul of the DNAse, incubating for 30 minutes, and repeating this procedure. RNA was immediately stored at −70 °C after inactivation of the DNAse before *ntrX* RT-PCR.

### RT-PCR of *ntrX*

The expression of the *ntrX* in the virulent (passage 7 and 11) and attenuated (passage 63 and 66) Senegal strains was determined using the primers *ntrX* qRT F1 (5′-GGAAAGATTGTATATTTCTG-3′) and *ntrX* qRT2 R1 (5′-ACCAGTAATGAGTATACGAC-3′) that amplify a 517 bp piece of the *ntrX* gene (Supplementary Figure 1). Amplifications were done using the OneStep RT-PCR kit (QIAgen, California, USA) with the following conditions: one Reverse transcriptase cycle at 50 °C for 30 minutes, a denaturating cycle at 95 °C for 15 minutes for activation of HotStart Taq, 35 cycles with a denaturating step 95 °C for 1 minute, an annealing step at 50 °C for 1 minute, and an amplification step at 72 °C for 1 minute, followed by an amplification cycle of 72 °C for 10 minutes. DNA from *E. ruminantium* Senegal passage 6 was used as positive control. Products were run in agarose gel and bands were visualized with SYBR safe.

## Results

### Virulent and attenuated Senegal strains genomic differences: SNPs and Indels

The variants found between the virulent and attenuated Senegal strains are shown in Table 1. We identified only two SNPs and three indels between the strains. The SNPs occur in *glyA*, a serine hydroxymethyltransferase involved in the interconversion of serine and glycine as well as tetrahydrofolate production, and a putative M16 protease. Both SNPs are nonsynonymous. The indels occur in a hypothetical gene, *map1-2*, a member of the *map1* family of outer membrane proteins, and *ntrX*, the response regulator of the putative nitrogen-sensing two component system. The hypothetical gene and *ntrX* both contain a 4bp deletion, while the *map1-2* gene contains a 2bp insertion. It is possible that the 4bp deletion in the *ntrX* duplicate arose due to the contraction of a 4mer sequence present in 2 copies in the *ntrX* gene (AGTTAGTT → AGTT) due to strand slippage during DNA replication. The *map1-2* gene appears to be a pseudogene in the virulent Senegal strain and the result of the insertion is to restore the open reading frame of the gene in the attenuated strain, making it a possible gain of function mutation. The hypothetical protein contains a Patatin domain and has sequence similarity to other patatin-like phospholipase family proteins that have lipolytic activity (Banerji and Flieger 2004).

**Table 1.** Variants between Senegal virulent and avirulent strains.

| Location (ERSA) | Gardel ortholog | Gene | Description | Mutation |
| --- | --- | --- | --- | --- |
| 299001 | ERGA_CDS_01780 |  | Hypothetical | Deletion 4bp |
| 1071844 | ERGA_CDS_06840 | *ntrX* | Response regulator for NtrY/X two-component system | Deletion 4bp |
| 1114557 | ERGA_CDS_07110 | *glyA* | Serine hydroxymethyltransferase | Non-Synonymous C388Y |
| 1217270 | ERGA_CDS_07720 |  | M16 family peptidase | Non-Synonymous S267L |
| 1426715 | ERGA_CDS_09130 | *Map1-2* | Map-family protein | Insertion 2bp |

### The Senegal *ntrX* gene is disrupted by segmental gene conversion from a nearby inverted partial *ntrX* duplication containing a 4bp deletion

A segment of the *ntrX* gene has been duplicated in all *E. ruminantium* genomes for which there is available genome sequence data (Figure 1A, Figure 2B). The duplicated section is inverted and covers 421bp close to the 5′ end of *ntrX* (not including the start codon), and it lies roughly 2kb downstream of the *ntrX* gene itself. The alignment between the *ntrX* and the duplicate region reveals that the 4bp deletion identified in the *ntrX* gene in the attenuated Senegal strain is also present in the duplicated segment in both the attenuated and virulent strains of Senegal, but not in any of the other strains (Figure 2A). The 4bp deletion in *ntrX* in the attenuated strain causes a frameshift which introduces a stop codon between the response regulator receiver and the sigma factor interaction domain regions (Figure 1B). There are also seven other mutations across the sequenced strains between *ntrX* and its duplicate that exhibit a pattern that is incongruent with the expected phylogeny (Figure 2A and 2B). At these residues, the *ntrX* gene and its duplicate are more similar within a strain than they are to their orthologous positions in the other strains, indicating recurrent gene conversion towards the 3′ end of the alignment in the different strains.

**Figure 1.**
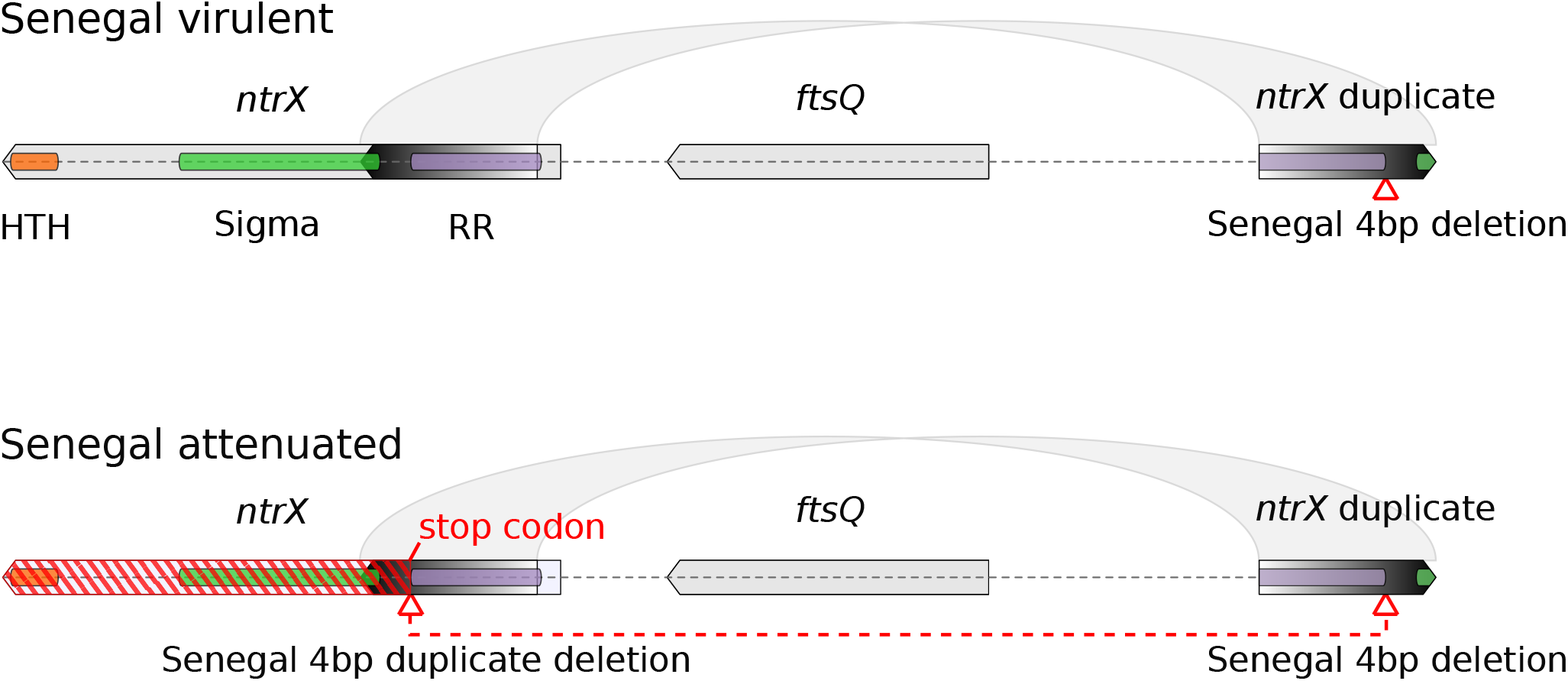
Structure of the *ntrX* gene in virulent and attenuated Senegal strains. The *ntrX* gene (left gene) is shown in its genomic context with its partial inverted duplicate (right gene) bounding the *ftsQ* gene in the virulent (top) and attenuated (bottom) strains. The inverted duplication is signified by the light grey arcs joining the two genes and the duplicated segment of the gene is depicted using a gradient from white to black. The encoded domains in *ntrX* are shown with orange (HTH: helix-turn-helix DNA-binding domain), green (Sigma: sigma factor-binding domain) and purple (RR: response regulator domain) bars within the *ntrX* gene. The 4bp deletion (red triangle) is found only in the *ntrX* duplicate in the virulent strain and in both the duplicate and the *ntrX* gene in the attenuated strain. The 4bp deletion causes a frameshift in *ntrX* in the attenuated strain creating a pseudogene (red hatching).

**Figure 2.**
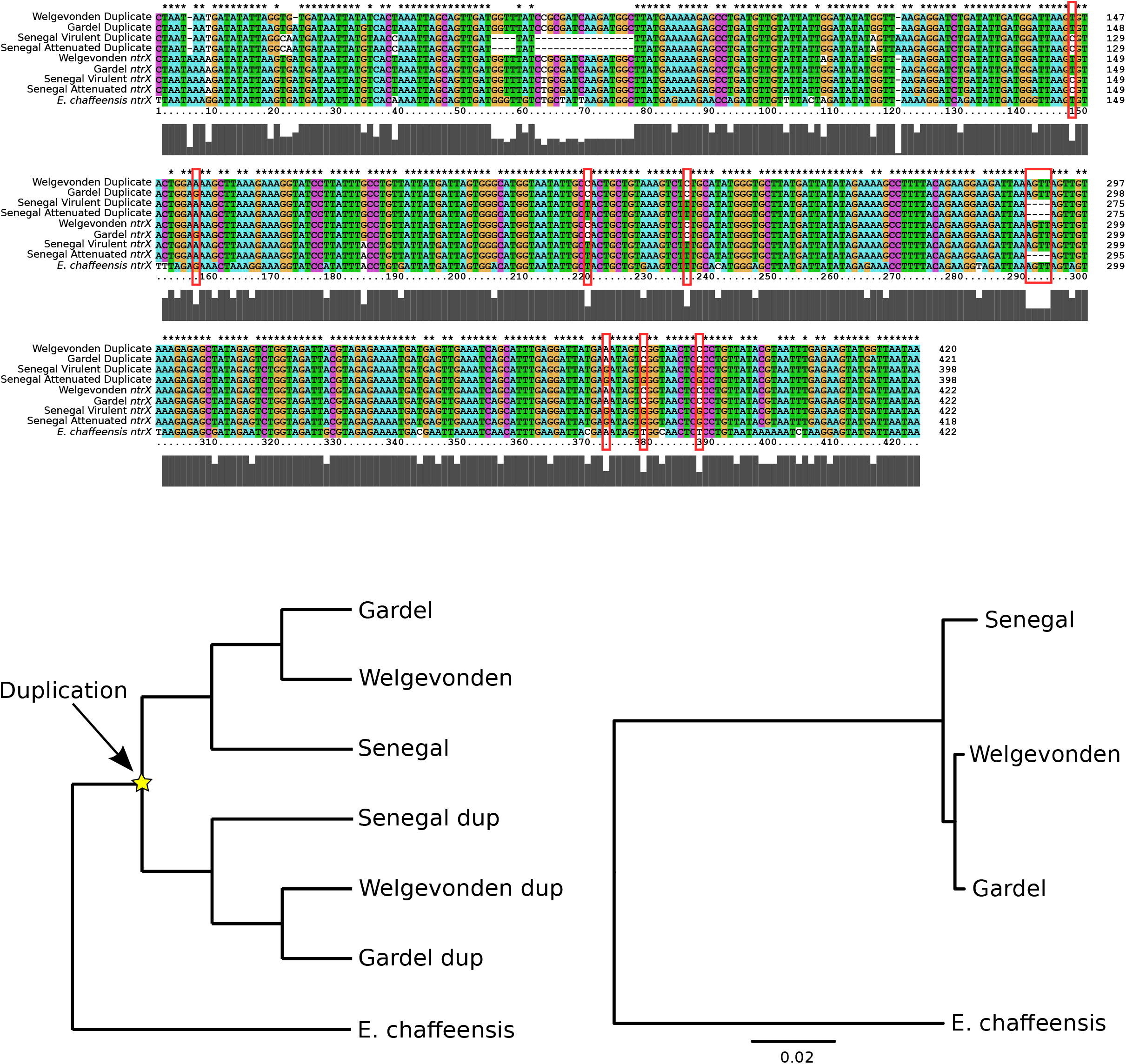
Evidence of gene conversion between *ntrX* and its inverted duplicate. A) An alignment showing the duplicated region (top four rows) of *ntrX* gene and the matching parental *ntrX* region (next four rows) in Welgevonden, Gardel and Senegal virulent and attenuated strains and in *E. chaffeensis* (last row) where there is no inverted duplicate. Alignment positions that show a closer relationship between *ntrX* and the duplicate within a strain than between strains, including the 4bp deletion in Senegal strain are bordered by red boxes. B) A phylogram (left) depicting the expected relationship between *ntrX* and its duplicate (yellow star) given that it is present in all *E. ruminantium* strains. The structure of the phylogram is based on the interstrain relationship reconstructed in the maximum likelihood phylogeny (right) by concatenating 54 orthologous ribosomal proteins from the strains and *E. chaffeensis*.

### NtrX is expressed in Senegal virulent strain but not in the attenuated strain

To confirm the knock out of *ntrX* expression in the attenuated strain, we performed a RT-PCR on RNA samples from the virulent (passage 7 and 11) and avirulent (passage 63 and 66) strains. RT-PCR resulted in the amplification of a 517 bp in one (Senegal passage 11 and not Senegal passage 7) of the two virulent samples (Supplementary Figure1, lane 5), confirming the expression of *ntrX*. No band was observed in either of the samples from the attenuated strain confirming that *ntrX* gene is not expressed in the attenuated Senegal strain (Supplementary Figure1, lanes 6 and 7).

## Discussion

### Candidate mutations for Senegal strain attenuation

The two nonsynonymous SNPs and the three indels identified between the virulent and attenuated Senegal strains provide a small number of candidates to explain the attenuation process in this strain. The *glyA* gene is involved in the interconversion of serine and glycine, and is necessary for virulence in *Salmonella Typhimurium* and *Brucella* species (Köhler et al. 2002; Xiang, Zheng, and He 2006; Jelsbak et al. 2014), however, the attenuation in *Brucella suis* was attained by Tn5 insertion to knock out the gene, and in the case of the SNP in our data, the protein is still probably produced. The substituted amino acid lies near the end of the protein (position 388/421) and outside of any predicted domains. An alignment of *glyA* in several Rickettsial species shows that while most of the protein is well conserved, the portion where the mutation occurs is variable across the species (Supplementary Figure 2), suggesting that the region is not vital for protein function. The *map1-2* gene is a member of a multigene family of Major Antigenic Proteins in the *Anaplasmataceae* (Pfam PF01617) (Dunning Hotopp et al. 2006), which are surface exposed and have been identified as potential vaccine targets (Bekker et al. 2002). Some members of the *map* gene family have been experimentally characterized as porins (Huang et al. 2007) and are suspected to be involved in host cell adhesion (Garcia-Garcia et al. 2004; Park, Choi, and Dumler 2003). Thus, its mutation appears to be a potential candidate for attenuation. However, a study of the *map1-2* gene in several different strains of *E. ruminantium* failed to provide evidence of transcription of this gene in any of the strains they tested (Senegal virulent and attenuated, Gardel, Welgevonden and Sankat 430) in either ticks or bovine endothelial cells (Bekker et al. 2002). A different study did however report the transcription of the *map1-2* gene in Welgevonden (van Heerden et al. 2004). The *map1-2* gene contains a 2bp deletion in the Senegal virulent strain (presumably rendering it non-functional) that is reverted in the attenuated strain. Curiously the deletion is not present in a previous study using a different isolate of the virulent Senegal strain (Bekker et al. 2005). Examination of the reads covering this indel in our data revealed that they fully support the deletion in the virulent Senegal, while it is not present in any reads in the attenuated strain. This result suggests that this deletion may be unique to the virulent strain isolate sequenced in this study. The lack of detectable transcription of the gene in some strains (Bekker et al. 2002) and the fact that the complete *map1-2* sequence is present in several other virulent strains of *E. ruminantium* including another Senegal isolate makes the reversion insertion unlikely to be the cause of attenuation of the Senegal strain.

The patatin domain-containing protein may play a role in the virulence of *E. ruminantium*, as patatin-like phospholipase proteins are putatively involved in host cell entry in Rickettsia (Rahman et al. 2010). Phospholipase proteins have been identified as virulence factors secreted by the Type III secretion system in *Pseudomonas aeruginosa* (ExoU), and the Type IV-B secretion system in Legionella pneumophilia (VipD) (Rahman et al. 2010). Bacterial pathogens also tend to contain more patatin-like-proteins than non-pathogens (Banerji and Flieger 2004), suggesting possible roles in virulence. However, their presence in non-pathogens also indicates that patatins are not always involved in virulence functions. The indel in the attenuated strain is close to, but outside of the predicted patatin domain, suggesting that the phospholipase activity might be maintained, but making the result of the indel unsure in terms of its effect on the function of the protein. Proteolysis can play roles in virulence at various levels in bacterial pathogens (Frees, Brøndsted, and Ingmer 2013), and although we could find no evidence for a known role of proteases from the M16 family in bacterial virulence, proteases from this family in *Toxoplasma gondii* have been suggested to play a possible role in host invasion by the parasite (Laliberté and Carruthers 2011).

The best attenuation candidate by far is the *ntrX* gene containing the 4bp deletion because it is the response regulator of one of only three two-component systems in *E. ruminantium* that are responsible for global regulation of various bacterial systems (Cheng et al. 2006; Kumagai et al. 2006; Cheng, Lin, and Rikihisa 2014), and it has been documented to affect virulence and survival in another mammalian pathogenic bacterium, *Neisseria Gonorrhoeae* (Atack et al. 2013). Two-component systems are involved in sensing of environmental or cellular signals and the downstream expression of genes, allowing bacteria to coordinate their gene expression in response to their environment or cellular state (Cheng et al. 2006; Jansen et al. 2015; Parkinson and Kofoid 1992; Nene and Kole 2008; Tanner et al. 2016). During infection, bacterial pathogens need to efficiently coordinate their metabolic activities with their virulence to allow for maximization of their growth and successful attack on the host cells, while avoiding host defences. Signals such as the availability of nutrients or metabolites may inform the bacteria of the optimal time to produce virulence factors, and the regulation of many bacterial virulence factors is linked with nutrient availability (Somerville and Proctor 2009; Barbier, Nicolas, and Letesson 2011). Thus, it is unsurprising that two-component systems have often been identified as virulence attenuators in various bacterial species given that they coordinate large and widespread changes in gene expression in response to environmental signals (Parkinson and Kofoid 1992; Cheng et al. 2006; Nene and Kole 2008; Jansen et al. 2015). NtrX lies at the crossroads of environmental sensing and gene expression and is therefore the ideal candidate to explain attenuation because perturbations to the NtrY/X system are likely to have major consequences for the growth, survival and the coordination of bacterial metabolism and virulence. Furthermore, *ntrX* as the attenuator explains the apparently biased nature of attenuation in the Senegal strain as compared to other *E. ruminantium* strains due to its genomic context, which will be discussed later.

### The attenuated strain has a possible growth advantage *in vitro*

The *in vitro* bovine endothelial cell medium offers a resource-rich environment for the bacteria to grow in without coming under attack from the host immune system. By performing multiple passages *in vitro*, the bacteria are placed under artificial selection to favour growth in cell culture. The *in vitro* adapted *E. ruminantium* strains are attenuated *in vivo* (Jongejan 1991; Pilet et al. 2012; Marcelino et al. 2015), suggesting that virulence is costly to the pathogen in terms of their reproduction when they are in a nutrient-rich environment free of the host’s active immune system. It has been noted that the attenuated Gardel strain exhibits a faster growth cycle that finishes 24 hours faster that the virulent strain (Marcelino et al. 2015). A faster growth rate may be a common feature of the attenuation process in *E. ruminantium* where passaging the bacteria in bovine endothelial cells *in vitro* places them under an artificial selection regime. Similarly, the attenuated Senegal strain has a faster growth cycle than the virulent strain (t-test, P=0.0003) (Supplementary Figure 3, Supplementary Table 1). It should be noted however that the virulent strain is much more variable in the time it takes and to lyse, an effect that could be due to a lack of synchronisation of infection in the cells with low loads of infectious bacteria or poor adaptation to culture. It has also been shown previously that the virulent Senegal strain reduces its lysis time as the passages progress, from 15-20 days at isolation to 6-12 days after growth in culture (Martinez et al. 1990), thus it is not possible to conclude that the faster growth of the attenuated strain in culture is directly due to the attenuation process, as it could be due to the adaptation to the cell culture conditions alone. The proteomics data from the Gardel strain exhibit a higher expression of proteins involved in cell division, metabolism, transport and protein processing for the attenuated strain compared to the virulent (Marcelino et al. 2015). Given the probable different mechanisms of attenuation (see below) between the strains, it is not certain that changes in metabolism in Senegal strain mirror those in Gardel strain. However due to the same selective environment of the *in vitro* media, it is certainly possible that the net results of attenuation on cellular metabolism are similar.

### Evidence for gene conversion in *ntrX*

The 4bp deletion in the partial inverted *ntrX* duplicate in the virulent Senegal strain is present in both the duplicate and the parent *ntrX* gene in the attenuated strain (Figure 1B), indicating a probable interaction and transfer of information between *ntrX* and its duplicate. Several other residues also evidence a transfer of genetic information between the duplicate regions, showing an incongruent relationship compared to that expected between the *ntrX* gene and its duplicate in *E. ruminantium* strains (Figure 2A). In the case of these residues, the *ntrX* gene and duplicate have the same residue within a strain, while it would be expected that the *ntrX* genes and duplicates should each be monophyletic, as the duplication event is ancestral to all available strains of *E. ruminantium*. The incongruent residues indicate a process of gene conversion occurring between *ntrX* and its duplicate that is ongoing independently in different strains. This process appears to be segmental with a tendency away from the 5′ of the duplicated region (with respect to the *ntrX* reading frame), as there is a deletion of several bases in the Senegal duplicate that has not been transferred to the parental *ntrX* gene, as well as a higher concentration of SNPs and indels towards the 5′ end (Figure 2A). Inverted repeats are prone to gene conversion processes because they can form cruciform DNA structures that bring the duplicate regions together as double-strands where mutations between them can be transferred due to one being used as a template for mismatch repair (Zhao et al. 2007).

There are some possible implications for the molecular evolution of the *ntrX* gene in *E. ruminantium* due to the inverted duplicate structure and segmental gene conversion. The partial duplicate is unlikely to be expressed, and mutations in the duplicate region are expected to be selectively neutral and unlikely to be weeded out by selection. Several mutations could potentially be transferred from the duplicate to *ntrX* at once, which could cause more drastic effects than the build up of single mutations. Although most mutations are expected to be either neutral or detrimental to gene function (T. Ohta 1992; Tomoko Ohta 2002; Kimura 1968), introducing several at once might provide an increased likelihood of selectively beneficial mutations, or combinations of mutations that may be beneficial in concert. Because mismatches in the cruciform structure formed by inverted duplicates could be resolved using either strand as a template i.e. gene conversion in both directions, the degeneration of the non-functional partial duplicate may be slowed.

### Evidence for a biased attenuation mechanism in the Senegal strain

Interestingly, the attenuation of the Senegal strain occurred originally in 11 passages (Jongejan 1991), and in at most 20 passages (Pilet et al. 2012) in the independently attenuated strain studied here (N. Vachiery, data not shown). In comparison, the Gardel strain required 230 passages to attenuate in a similar manner (Marcelino et al. 2015), and the Welgevonden strain grown in the same culture was still virulent at 231 passages (Zweygarth et al. 2005) (it was only through passaging in canine macrophages that Welgevonden attenuation was attained). The large difference in time required for attenuation between Senegal and the other two strains suggests that there are probably different mechanisms of attenuation in these strains, while the similarity in the number of passages required to independently attenuate virulent Senegal strains suggests that they probably attenuated by the same biased mechanism. The transfer of a 4bp deletion from the *ntrX* partial inverted duplicate to the parental gene by gene conversion (Figure 1) offers a simple explanation for how the Senegal strain attenuates quickly compared to the other strains where the duplicate does not contain a deletion, and it is consistent with the evidence of ongoing gene conversion at these loci in the different strains. The initial 4bp deletion in the duplicate was presumably neutral and stochastic, but once in place allows for a “loaded die” mechanism of virulence attenuation in the Senegal strain that is not revertible by gene conversion. The attenuation of the other strains presumably relies on a different mechanism, i.e. the accumulation of random mutations in the genome until attenuation emerges by the disruption of a system that is vital *in vivo* but dispensable *in vitro*. The fact that the Senegal strain has been rapidly attenuated on two separate occasions suggests that *ntrX* pseudogenisation is selectively advantageous *in vitro*, and that NtrX regulates systems that are necessary for infection, but that are dispensable in an environment containing readily available nutrients and no need to circumvent the host cell defences.

### The NtrY/X 2CS

While the *NtrY/X* system is widely distributed across bacterial species, it appears that its function is not well conserved. The *NtrY/X* 2CS was first identified and characterised in the free-living nitrogen fixing bacterium *Azorhizobium caulinodans* ORS571 where it was found to be involved in the sensing, metabolism and fixation of nitrogen and nitrogen-related compounds (Pawlowski, Klosse, and de Bruijn 1991). The *ntrY* gene was identified as a probable transmembrane sensor kinase protein in *A. caulinodans* and *A. brasilense*, while *ntrX* is the response regulator that modulates downstream gene expression (Kumagai et al. 2006). A role in the regulation of respiratory enzymes including the cytochrome c oxidase unit has been identified in the human pathogenic beta-proteobacterium *Neisseria gonorrhoeae* (where there is no evidence that *ntrX* modulates the expression of nitrogen metabolism genes) and the free-living alpha-proteobacterium *Rhodobacter capsulatus*, where it is also involved in the expression of photosynthesis genes (Atack et al. 2013; Gregor et al. 2007). The *N. gonorrhoeae ntrX* mutants exhibit a reduced ability to invade and survive in human cervical epithelial cells, suggesting that *ntrX* may play a role related to pathogenicity in this organism (Atack et al. 2013). In the animal pathogenic alpha-proteobacterial *Brucella* spp, *ntrY* has been identified as an attenuation gene by transposon mutagenesis (Foulongne et al. 2000), and the *NtrY/X* system has also been implicated as a redox sensor and regulator (Carrica et al. 2012). In these species, *ntrY* contains a haem-binding PAS domain that is involved in the sensing of oxygen tension in the cell, an important function in a bacterium that needs to adapt to host cell defences such as reactive oxygen and nitrogen species (Carrica et al. 2012). Control of the respiratory gene expression may enable the adaptation and survival of intracellular oxidase-positive bacteria (Atack et al. 2013), and could relate to their central metabolism and the coordination of their growth and production of virulence factors.

The *NtrY/X* system is present in most sequenced bacteria of the order Rickettsiales, with notable absences in *Wolbachia* species and *Neorickettsia sennetsu* (Cheng et al. 2006). There are no studies that characterise the *NtrY/X* system in *E. ruminantium*, although some studies have been performed in closely related *E. chaffeensis* and *Anaplasma* species (Cheng et al. 2006; Cheng, Lin, and Rikihisa 2014; Kumagai et al. 2006), which due to their evolutionary proximity to *E. ruminantium*, represent the best available models of the system for our work. In these species, the *NtrY/X* system represents one of only three identified 2CSs (Cheng et al. 2006; Kumagai et al. 2006; Cheng, Lin, and Rikihisa 2014). Unlike most bacterial systems examined where 2CS sensor kinases tend to be membrane-bound, in these species (and also in *E. ruminantium*) *ntrY* does not contain a putative cleavage signal peptide, and is thus predicted to be a non-membrane protein (Kumagai et al. 2006; Cheng, Lin, and Rikihisa 2014; Cheng et al. 2006). In confirmation, it was detected in the soluble fraction of *E. chaffeensis* fractionation (Kumagai et al. 2006; Cheng, Lin, and Rikihisa 2014), indicating that it is probably a sensor of internal cellular environmental signals (Cheng, Lin, and Rikihisa 2014). In *E. chaffeensis*, a PAS domain is not present in *ntrY* (Cheng, Lin, and Rikihisa 2014), also indicating a likely different sensory function to those species where a PAS domain is present. The *ntrX* gene in *E. chaffeensis* and *E. ruminantium* contains a predicted sigma-54 (rpoN/sigmaN) binding domain, but puzzlingly the only identified sigma factors in this organism are sigma-70 and sigma-32, indicating that intracellular development is regulated by constitutive sigma-70 promoters (Kumagai et al. 2006; Dunning Hotopp et al. 2006; H. Liu et al. 2013; Rikihisa 2015).

In *Ehrlichia chaffeensis*, the three present 2CSs are expressed sequentially at different times during the cell cycle with slight overlap (Rikihisa 2015), implicating each in the regulation of different parts of the growth cycle. The *ntrX* gene is expressed at the end of the lag phase and during early exponential growth, coinciding with the transition in cellular morphology from dense-cored cells to reticulate cells (Rikihisa 2015), suggesting that *ntrX* may coordinate the initiation of exponential growth and/or changes in cellular morphology and may be involved with coordinating invasion of the host cell. Interestingly, the *virB/D* genes that make up the Type IV Secretion System (T4SS) implicated in bacterial virulence and the immunogenic P28 outer membrane protein (ortholog of Map1 in *E. ruminantium*) implicated in nutrient uptake are also upregulated during exponential growth (Cheng, Wang, and Rikihisa 2008; Kumagai, Huang, and Rikihisa 2008). Thus, it is likely that virulence factor expression and nutrient uptake from host cells may occur during this phase of the cell cycle when the bacteria need to acquire nutrients from their hosts to allow maximum growth. Of the 2CSs in Ehrlichia, only NtrX and CtrA have response regulators with DNA-binding and transcriptional regulation activities (Rikihisa 2015), and as they each are expressed at different points in the cell cycle it is likely that the regulon of NtrX is large like that of CtrA (Cheng et al. 2011).

In *Ehrlichia chaffeensis* two genes, *putA* and *glnA*, involved in the conversion of proline to glutamate and glutamate to glutamine, are simultaneously upregulated during early cell growth by NtrX, which directly binds their promoters in its phosphorylated form in the presence of proline and glutamine (Cheng, Lin, and Rikihisa 2014). Proline and glutamate are likely to be the primary carbon sources in *E. ruminantium* based on the presence of transporters in the genome (Collins et al. 2005), although glutamine uptake has also been demonstrated by the P28 porin in *E. chaffeensis* (ortholog of Map1 in *E. ruminantium*) (Cheng, Lin, and Rikihisa 2014; Kumagai, Huang, and Rikihisa 2008). Biosynthesis of nitrogenous compounds like DNA, RNA and proteins also utilise glutamine and glutamate (Cheng, Lin, and Rikihisa 2014). It has been demonstrated that proline and glutamine are critical for *E. chaffeensis* host cell invasion and proliferation, and that pretreatment of the cells with these two amino acids enhances bacterial growth and shortens the lag phase of the growth curve (Cheng, Lin, and Rikihisa 2014). NtrX has a direct interaction with *putA* and *glnA* promoters, so the NtrY/X system probably plays a vital role in sensing and responding to the availability of these and possibly other nutrients in *Ehrlichia* species. To coordinate metabolism and virulence, cues like nutrient availability are vital, so removing or perturbing these signals are likely to disrupt the normal growth-cycle progression. Blocking proline and glutamine inhibits cell growth in *E. chaffeensis*, at least in part because they degrade CtrA (Cheng et al. 2006; Cheng, Lin, and Rikihisa 2014). CtrA binds the origin of replication to inhibit DNA replication (Cheng, Lin, and Rikihisa 2014), and is predicted to be important in cell cycle progression and morphology (Cheng et al. 2011).

We propose two mechamisms for how *ntrX* pseudogenisation could affect the growth of the Senegal strain *in vivo*. Firstly, a possible outcome of *ntrX* knockout is an increased concentration of proline and glutamine in the cell due to the lack of signal from NtrX-P upregulating *putA* and *glnA*, thus increasing the degradation of CtrA bound to the origin and initiating DNA replication. In this way, NtrX could contribute to cell cycle progression. Secondly, the lack of NtrX signal might erroneously signal to the cell that it is in nutrient-limited conditions, i.e. conditions similar to the end of exponential growth, or that it has reached the end of the *ntrX* expression period, causing the induction (directly or through an indirect signal) of the expression of *pleC/D*, which usually regulates the growth-cycle after NtrX (Rikihisa 2015). This could effectively cause the cells to skip from lag phase to mid-exponential growth, causing a truncation of the growth cycle. Further work is required to validate or disprove these different hypotheses.

The lack of *ntrX* expression in the attenuated Senegal strain shows that NtrX is not vital for *E. ruminantium* cell survival. The fact that the *ntrX* pseudogenisation is tolerated (and perhaps selected for) *in vitro* but attenuates the bacterium *in vivo* indicates that the NtrX regulon contains genes involved in pathogenic growth and survival that are indispensable for *in vivo* infection, but may also present a production cost to the bacterium. The possible selection acting to attenuate the bacterial cells could be acting to increase growth rate by reducing the production of superfluous proteins in the *in vitro* environment.

## Conclusion

The best candidate mutation by far to explain the attenuation in *E. ruminantium* Senegal is the 4 bp deletion found in the *ntrX* gene. As a global regulator expressed during the invasion of host cells (late lag-phase and early exponential growth) that is a part of a two component system used for modulating expression of genes in relation to some environmental signal, it is an ideal candidate to explain the virulence attenuation of the Senegal strain. The presence of common mutations between *ntrX* and a nearby inverted partial duplicate that are found within different strains provides evidence of gene conversion, and explain how the 4bp deletion arose in the *ntrX* gene in the attenuated Senegal strain where the duplicate already contains the 4bp deletion. This mechanism explains the fast attenuation of the Senegal strain on two independent occasions when passaged in an *in vitro* environment compared to other attenuated strains, and at the same time suggests a growth advantage in the *in vitro* medium for the attenuated strains. This is the first study to outline the probable genetic basis for virulence attenuation in *Ehrlichia ruminantium* and provides a promising mutational target to speed up vaccine development in other strains.

## Acknowledgements and funding information

The authors would like to thank Ludovic Pruneau for technical assistance with growth of the Senegal strains and Valérie Barbe at the French National Sequencing Centre Genoscope for sequencing. The genome assemblies for the ERSA and ERSB strains have been submitted to Genbank (Accessions GCA_002019755.1 & GCA_002189655.1, respectively). This work was supported by European project “EPIGENESIS” FP7-REGPOT-2012-2013-1 (31598).

