## Supplemental Figures and Table for "Possible biased virulence attenuation in the Senegal strain of *Ehrlichia ruminantium* by *ntrX* gene conversion from an inverted segmental duplication"

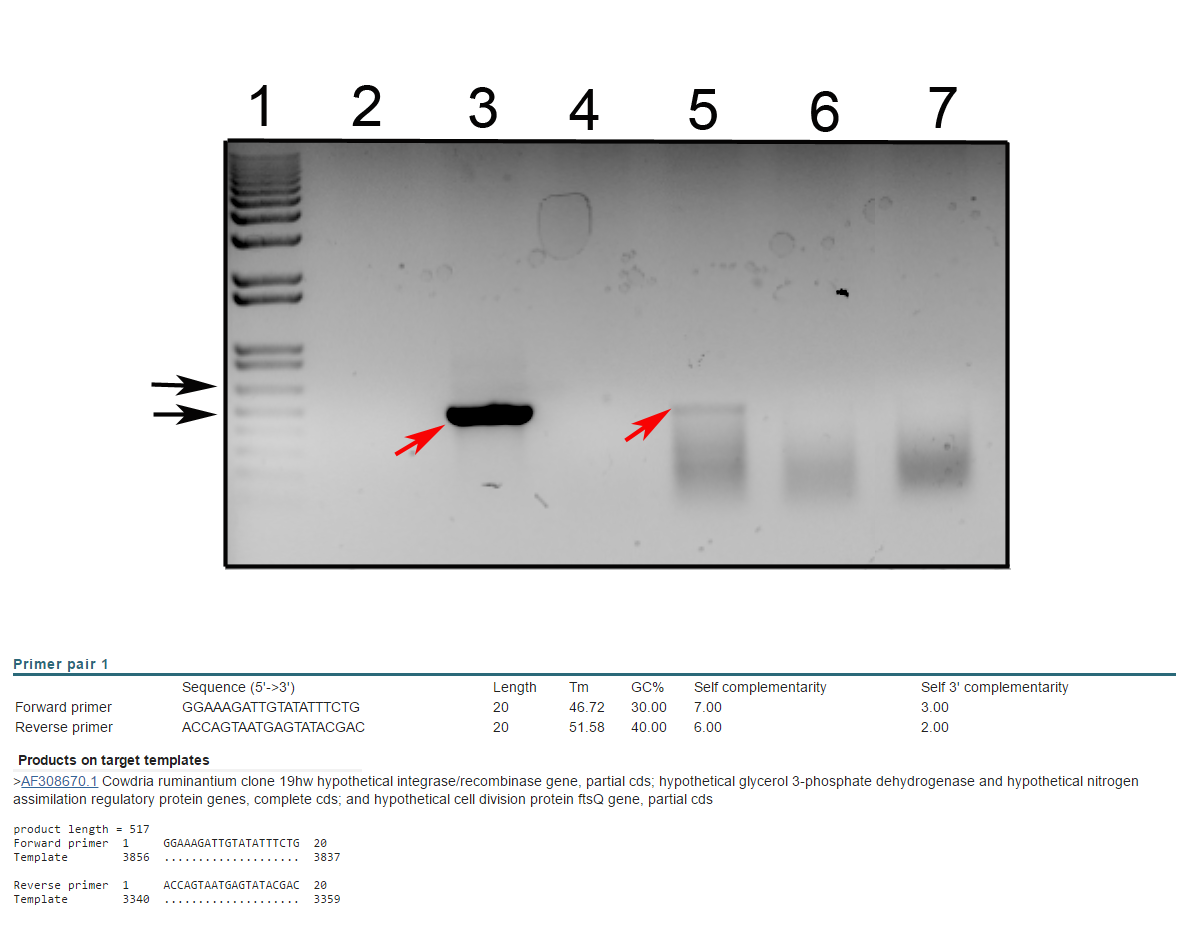

**Supplementary Figure 1. RT-PCR amplification of *ntrX* in the virulent and attenuated strains**

The expression of the *ntrX* gene was analysed on total RNA samples from the virulent and attenuated strains of *E. ruminantium* Senegal. RNA samples were prepared from infected cell layers with 80 – 90% infection. RT-PCR amplification of was performed using water as negative control (lane 2), DNA from passage 6 as positive control (lane 3), passages 7 (lane 4) and 11 (lane 5) of the virulent Senegal strain and in passages 63 (lane 6) and 66 (lane 7). Expression of the gene was confirmed in passage 11 by amplification of the 517 bp band, which was also observed in the positive control (red arrows). The size of the band was confirmed by size comparison with the 1 kb plus ladder (lane 1; black arrows show 500 and 650 bp bands). No RNA amplification was observed for lane 4 probably because no RNA was present.

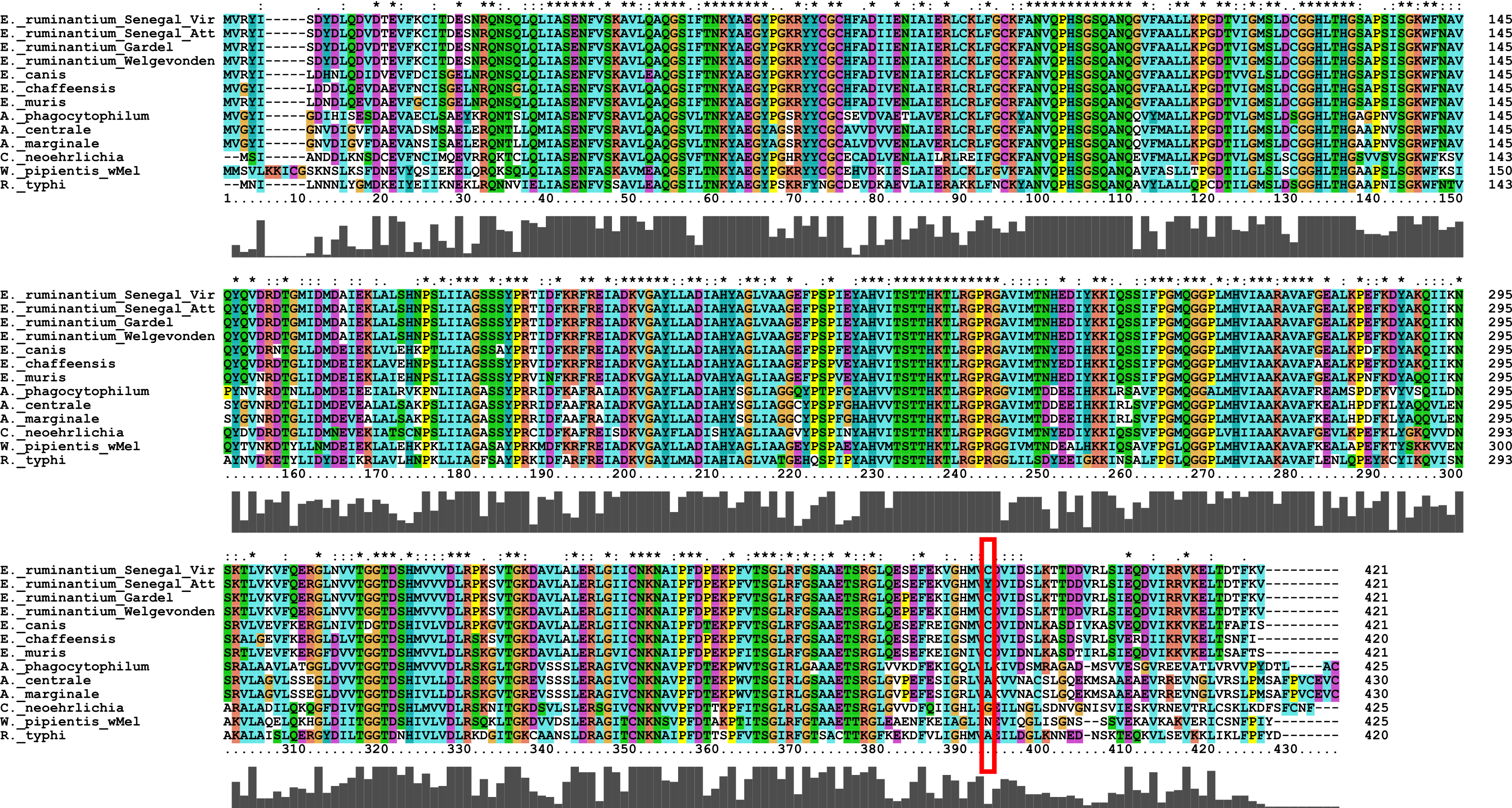

**Supplementary Figure 2. Alignment of *glyA* in Rickettsial species.**

The alignment includes the *glyA* gene from a range of related Rickettsial species and the nonsynonymous mutation in the Senegal strain is outlined with a red rectangle. The region in which the mutation is located is not well conserved across the species, as shown by the histogram below the alignment. The genes were identified using BLAST (Altschul et al. 1990) against the Gardel strain *glyA* gene and were aligned using Muscle (Edgar 2004) and the image was output using ClustalX (Larkin et al. 2007).

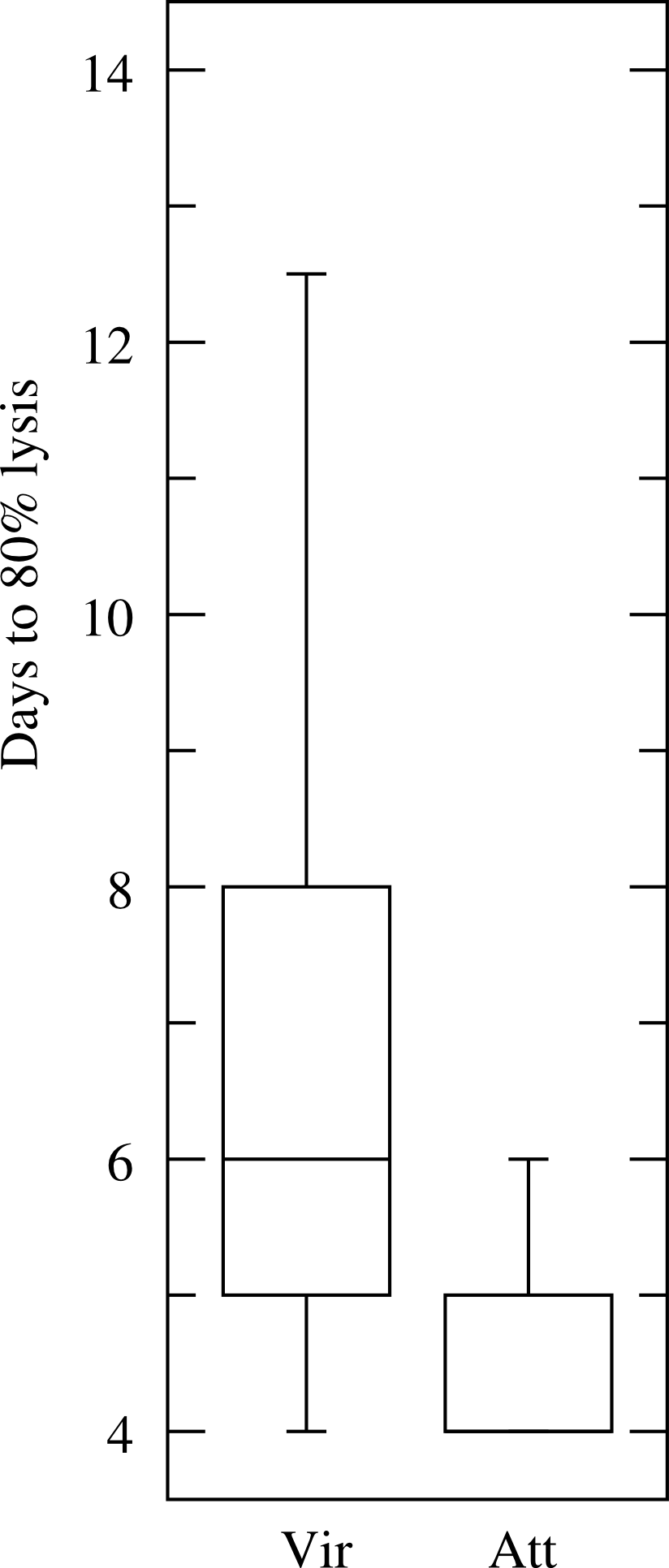

**Supplementary Figure 3. Boxplot of lysis times (days) for virulent and attenuated Senegal strains**

The attenuated strain shows significantly faster lysis times (t-test, P=0.0003). The boxplot was created at (<http://www.physics.csbsju.edu/stats/t-test_bulk_form.html>) using the data provided in Supplementary Table 1.

**Supplementary Table 1. Days for lysis of virulent and attenuated Senegal strain passages.**

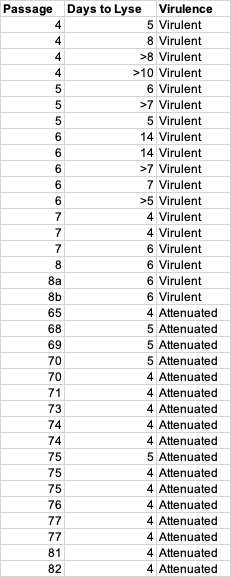
